# Spectral Characterization of a Blue Light-Emitting Micro-LED Platform and Microbial Chromophores for Therapeutic Applications in Skin Conditions

**DOI:** 10.1101/2024.03.05.582921

**Authors:** Hannah J. Serrage, Charlotte J. Eling, Pedro U. Alves, Andrew J. Mcbain, Catherine O’neill, Nicolas Laurand

## Abstract

The therapeutic application of blue light (380 – 500nm) has garnered considerable attention in recent years as it offers a non-invasive approach for the management of prevalent skin conditions including acne vulgaris and atopic dermatitis. These conditions are often characterised by an imbalance in the microbial communities that colonise our skin, termed the skin microbiome. In conditions including acne vulgaris, blue light is thought to address this imbalance through the selective photoexcitation of microbial species expressing wavelength-specific chromophores, differentially affecting skin commensals and thus altering the relative species composition. However, the abundance and diversity of these chromophores across the skin microbiota remains poorly understood. Similarly, devices utilised for studies are often bulky and poorly characterised which if translated to therapy could result in reduced patient compliance. Here, we present a clinically viable micro-LED illumination platform with peak emission 450 nm (17 nm FWHM) and adjustable irradiance output to a maximum 0.55±0.01 W/cm^2^, dependent upon the concentration of titanium dioxide nanoparticles applied to an accompanying flexible light extraction substrate. Utilising spectrometry approaches, we characterised the abundance of prospective blue light chromophores across skin commensal bacteria isolated from healthy volunteers. Of the strains surveyed 62.5% exhibited absorption peaks within the blue light spectrum, evidencing expression of carotenoid pigments (18.8%, 420 – 483 nm; *Micrococcus luteus, Kocuria* spp.), porphyrins (12.5%, 402 – 413 nm; *Cutibacterium* spp.) and potential flavins (31.2%, 420 – 425 nm; *Staphylococcus* and *Dermacoccus* spp.). We also present evidence of the capacity of these species to diminish irradiance output when combined with the micro-LED platform and in turn how exposure to low-dose blue light causes shifts in observed absorbance spectra peaks. Collectively these findings highlight a crucial deficit in understanding how microbial chromophores might shape response to blue light and in turn evidence of a micro-LED illumination platform with potential for clinical applications.

## 1 Introduction

Skin conditions including atopic dermatitis and acne vulgairs not only dramatically impact upon patient’s quality of life but also present an enormous financial burden on health services (1). Current indications suggest that manipulating interactions of the skin microbiome (the collection of microbes that colonise our skin) could prove a promising approach for the management of skin conditions. Where certain microbes that colonise our skin are often overrepresented in disease (2).

Blue light (400 – 500 nm) presents a non-invasive approach for the management of skin conditions and has proven effective for conditions such as acne vulgaris and atopic dermatitis (3). Acne patients harbour a microbiome often dominated by strains of *Cutibacterium acnes*. These strains tend to produce higher levels of porphyrins, heterocyclic compounds that exhibit a soret band at 405 nm (4). The efficacy of blue light is thus often attributed to the photo-excitation of porphyrins, resulting in the production of cytotoxic reactive oxygen species (ROS) and a reduced abundance of *Cutibacterium acnes* (5).

Comparatively atopic dermatitis (AD) patients often experience flares, characterised by an elevated abundance of *Staphylococcus aureus* (6). *S. aureus* strains frequently express a golden carotenoid pigment with absorbance peaks at 440, 462 and 491 nm (7). However, despite a handful of studies demonstrating the efficacy of blue light for the management of atopic dermatitis (8), whether this results from photo-excitation of these endogenous carotenoid pigment and a decreased abundance in *S. aureus* is yet to be seen. Where current evidence remains conflicting, with some studies suggesting the antioxidant properties of carotenoid pigments may in fact enhance survival in response to light (9, 10), whilst others present evidence of the bactericidal properties of blue light against *S. aureus* (11, 12)

Despite these conditions being characterised by elevated levels of certain species and strains, the microbiota in these conditions remains relatively diverse (13). With commensal *Staphylococcus* spp. (particularly *Staphylococcus hominis* and *Staphylococcus epidermidis*) and *Corynebacterium* spp. remaining abundant during AD flares (6). Skin commensals that predominate within the microbiome express a host of pigments, proteins and molecules that may prove capable of eliciting a response to blue light, resulting in a shift in microbiome composition and subsequent modulation of inflammation characteristic of disease flares (14, 15). These photosensitive targets span flavins (16), porphyrins (17) and pigments including carotenoids (18) and phenazines (19) each with characteristic spectral properties. Photoexcitation of such targets has proven to exert profound effects on microbial behavior. For example, *Acientobacter baumannii*, an opportunistic nosocomial pathogen implicated in soft tissue infections expresses a blue-light sensing using flavin (BLUF) domain whose excitation at 470 nm modulates biofilm formation, virulence and bacterial motility (20-23). These properties are not limited to environmentally acquired pathogens and skin commensals *Micrococcus luteus* and *Kocuria arsenatis* express carotenoid pigments that act as antioxidants conferring resistance against ultraviolet radiation exposure (24, 25). However, the diversity and abundance of such pigments and proteins across the skin microbiome remain poorly characterised.

Another key deficit in studies assessing the prospective efficacy of blue light in the management of skin conditions is the devices utilised, which are often bulky and poorly characterised. Parameters applied such as wavelength (nm), irradiance (mW/cm^2^), and dose (J/cm^2^) are often misreported or not reported at all rendering it difficult to draw conclusions surrounding the use and mechanism of action of blue light in the management of skin conditions (26). Concomitantly, user design and functionality are rarely coupled to *in vitro* experimentation meaning the devices used are not clinically translatable. As light technology has significantly advanced in recent years, the application of flexible and portable devices for the management of prevalent skin conditions offers a new frontier in light therapy (27-32).

In this study, we demonstrate the thorough spectral characterisation of a light, optionally flexible, blue light emitting (450nm) micro-LED (μLED) illumination platform, which presents an attractive device for the management of skin conditions (31). Utilising this robust and reproducible system to deliver homogeneous irradiance on a desired location, we show that bacterial species abundant across the acne, AD and healthy skin microbiomes exhibit differential capacities to attenuate light output from the μLED array and map capacity to absorbance spectrum peaks within the range of 440 – 450 nm. We also show how light exposure alters the profile of absorbance spectra, indicating potential photoexcitation of pigments: providing insight into prospective chromophores expressed across the skin microbiome, and in turn how we might exploit these in the management of skin conditions using blue light.

## 2 Methods

### μLED platform

The platform comprised a blue emitting, 1 by 10 array of flip-chip GaN-based μLEDs with emission through their sapphire substrate (300 μm thick). The emitting area of each μLED was 100 x100 μm^2^ and the separation between μLEDs encompassing the array was 720 μm. Because of diffraction, the feature size of a μLED emission increases from 100 μm to∼ 300 μm at the sapphire output facet (29). The μLEDs were connected in parallel and the array was mounted on a printed circuit board (PCB). In this study, the μLED array was driven at 120 mA (2.94 V) giving an optical power of 10 mW. Details of array design and characterisation can be found elsewhere (30). The μLED array was coupled to a 1 mm x 75 mm x 25 mm glass slide, which guided light within by total internal reflection in an edge-lit configuration, the size of the μLED emission area at the sapphire facet (smaller than the thickness of the slide) ensuring maximisation of the coupling efficiency (Fig. 1). A secondary (scatter) substrate was added on top of the guiding layer to extract the guided light, illuminating the sample above (Figure 1A). The scatter substrates were fabricated from a ratio of 5 to 1 silicone (polydimethylsiloxane; PDMS, Sigma Aldrich) to cross-linker (RTV615), doped with increasing quantities of Ti(V) oxide anatase nanoparticles (TiO_2_, Sigma, US) nanoparticles (0, 0.05, 0.1, 0.25, 0.5, 0.75, 1 and 2% w/v) and left to set at room temperature for 48 hours to ascertain optimal concentration required to potentiate irradiance output.

**Figure 1:**
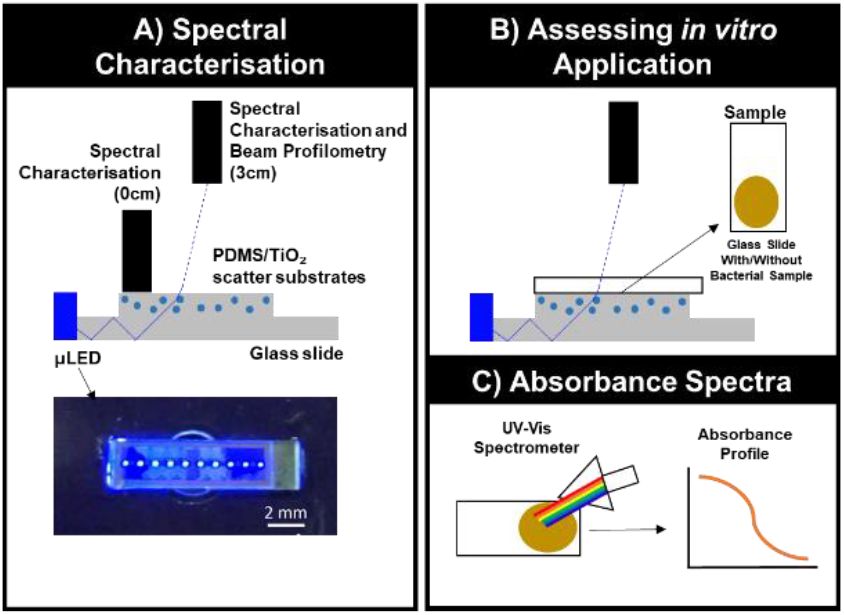
Diagrammatic Representation of Spectral Characterisation and in vitro Application. **A)** Setup utilised to characterise how different concentrations of TiO_2_ potentiate irradiance output and variability in irradiance output across the substrate surface, measured in direct contact (0cm) with surface and compared to values acquired 3cm above. **B)** Glass slides ± bacterial samples were placed directly on top of scatter substrates and changes in irradiance/wavelength output recorded 3 cm from the substrate surface. **C)** Absorbance profiles were collected using a UV-Vis spectrometer to determine whether expression of microbial chromophores (denoted by peaks in absorbance profiles) could be used to predict impact on irradiance output from light source.

### Spectral Characterisation

A UV-Vis Spectrometer (Avantes Starline AvaSpec-2048L, The Netherlands) coupled to a 200 μm-core optical fibre, with a 0.7 nm spectral resolution and calibrated to National Institute of Standards and Technology (NIST) standards was employed to assess spectral irradiance and wavelength delivery through 15 x 15 mm^2^ PDMS scatter substrates of the μLED array illumination platform containing increasing concentrations of TiO_2_ from 0 – 2% w/v. Spectral characteristics were recorded at 3 cm above or mapped in direct contact with substrates or samples (Figure 1A-B). Absolute irradiance was determined from the integral of the spectral irradiance (260 – 740 nm).

### 2D Irradiance Profiling

An N-BK7 Plano-convex lens and neutral density filter were positioned 9 cm and 16 cm from a CMOS camera (Thorlabs, US) to image the top surface of the LED array platform (Figure 2). Images were acquired using imaging software (Thorcam). 2D irradiance mapping was visualised using ImageJ (NIHR, US) and using a spectrometer as described above (Spectral Characterisation) for pre-calibration.

**Figure 2:**
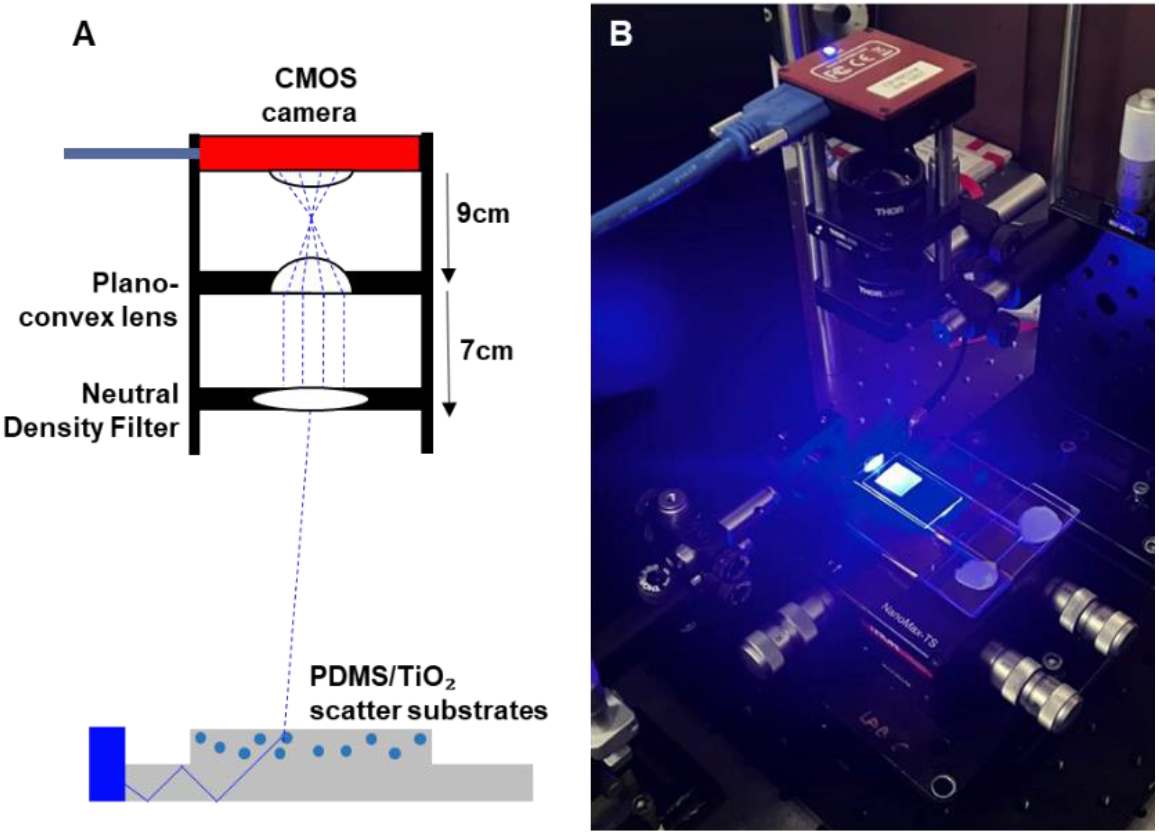
(A) Diagram and (B) Images captured of micro-LED array coupled to 0.5% TiO_2_ scatter substrate via Thorcam imaging software.

### Bacterial Isolation, Culture and Preparation

Strains used in this study were isolated from healthy volunteers (study approved by University of Manchester Ethics committee; 2019-6208-10419). In brief, swabs were dipped into pre-warmed phosphate buffered saline (PBS) and samples collected in duplicate into phosphate buffer (20mM Na_2_HPO_4_, KH_2_PO_4_, 0.1% Tween 80, 0.03% cysteine-hydrochloride, pH 6.8 (all Sigma-Aldrich)) from the volar forearm, forehead and scalp. Samples were spread onto fastidious anaerobe agar + 5% horse blood (Fisher Scientific), Wilkins-Chalgren anaerobe agar (WCA, Thermo Scientific) ± 0.025/0.1 % Tween 80 or Tryptic Soy agar (TSA, Oxoid)) ± 0.1 % Tween 80 and incubated for < 7 days under aerobic (37°C) or anaerobic conditions. Subsequently, colonies of distinct morphology were subbed onto WCA/TSA ± 0.1 % Tween 80 and once purity was reached, frozen in brain heart infusion + 0.5% yeast and 30% glycerol (Fisher Scientific).

To ascertain identity single colonies were picked from agar plates, and placed into a PCR reaction mix and DNA released through heating at 95°C for 10 minutes and the 16s region amplified (F; 5’-TGTAAAACGACGGCCAGT-3’, R; 5’-CAGGAAACAGCTATGACC-3’) via standard thermocycling. PCR products were confirmed on 0.8% agarose gel at 90V for 60 min and purified using Qiaquick PCR purification kit according to the manufacturer’s instructions (Qiagen). DNA concentration and quality were assessed, resuspended in nuclease-free water with forward primer and samples submitted to the genomic technologies core facility at the University of Manchester. Species identity was then assessed via input of sequences to nucleotide BLAST (https://blast.ncbi.nlm.nih.gov/). The strains and growth conditions employed in this study are listed in Table 1. Following the initial culture, colonies were inoculated upon glass microscopy slides, air-dried and fixed via heat.

**Table 1:** Species, strains and culture conditions used in this study.

| Species | Strain | % identity | Media | Incubation (h) | Conditions |
| --- | --- | --- | --- | --- | --- |
| *Corynebacterium afermentans* | CIP103499 | 93 | TSB + 0.1% Tween 80 | 16 | Aerobic |
| *Corynebacterium tuberculostearicum* | Medalle X | 100 | TSB + 0.1% Tween 80 | 16 | Aerobic |
| *Cutibacterium acnes* | JCM 6425 | 98 | WCB | 48 | Anaerobic |
| *Cutibacterium acnes* (177) | HKG366 | 98 | WCB | 48 | Anaerobic |
| *Cutibacterium gransulosum* | NCTC 11865 | 96 | WCB | 48 | Anaerobic |
| *Cutibacterium namneste* | NTS 31307302 | 97 | WCB | 48 | Anaerobic |
| *Dermacoccus nishinomiyaensis* | DSM 20448 | 98 | TSB | 16 | Aerobic |
| *Kocuria arsenatis* | CM1E1 | 99 | TSB | 16 | Aerobic |
| *Kocuria rhizophila* | TA68 | 97 | TSB | 16 | Aerobic |
| *Micrococcus luteus* | NCTC 2665 | 100 | TSB | 16 | Aerobic |
| *Staphylococcus aureus* | S33R | 98 | TSB | 16 | Aerobic |
| *Staphylococcus aureus* (wound) | NCTC 6571 | 98 | TSB | 16 | Aerobic |
| *Staphylococcus capitis* | ATCC 27840 | 97 | TSB | 16 | Aerobic |
| *Staphylococcus epidermidis* | NBRC 100911 | 98 | WCB | 16 | Aerobic |
| *Staphylococcus hominis* | GTC 1228 | 98 | TSB | 16 | Aerobic |

### Effects of Bacteria on μLED Platform Irradiance and Spectral Output

Slides comprising bacteria and standards were placed directly upon a 0.5% TiO_2_ scatter substrate, and changes in irradiance and wavelength output were recorded via spectrometer at a distance 3 cm from the substrate (Fig 1B).

### Absorbance Profile Measurements

Absorbance profiles of bacterial species inoculated upon glass slides (Fig 1C) were acquired using a UV-Vis spectrometer (ThermoFisher Scientific, US) pre– and post-exposure to the μLED platform blue light (λmax: 450 nm, FWHM: 17nm, 2.76 J.cm^2^) relative to a reference glass slide. Once acquired, absorbance profiles were plotted between 380 nm – 520 nm.

### Statistical Analyses

Data were processed utilising Excel software (Microsoft) and analyses were performed using Prism (GraphPad Software, California, US). All experiments were performed at least in duplicate unless otherwise stated, and data were analysed using a general linear model (GLM) followed by one-way ANOVA and Tukey/Dunnett’s test.

## 3 Results

### TiO_2_ scatter substrates potentiate irradiance output from the μLED array platform

TiO2 nanoparticles embedded within PDMS substrates scatter the μLED light guided in the underlying glass slide (Figure 1A). This ensures a homogenous irradiance over the area where the samples are placed (Figure 1B). Spectral characterisation utilising the set-up described in Figure 1, revealed the μLED array emitted a peak wavelength of 450.90 ± 0.35 nm (Figure 3A). When irradiance measurements were acquired 3 cm from the substrate, a 0.5% TiO_2_ scatter substrate elevated maximal irradiance to a peak 0.55 mW/cm^2^ (Figure 3B), but had no significant impact on wavelength output (Figure 3C). In which variability in the PDMS substrates containing no TiO_2_ (0% w/v) was due to a low detectable signal. Output variability significantly reduced following the application of even low concentrations of TiO_2_ (0.05 % w/v, 449.05 ± 0.35 nm) due to reduced confounding noise within the spectrometer range (216.2 – 740.074nm) as an increased irradiance signal was detected (complete range not plotted Figure 3A). 2D irradiance mapping indicated average irradiance across the 0.5% w/v TiO_2_ substrate varied from 1.33 mW/cm^2^ in areas closest to the μLED array coupling plane compared to 0.31 mW/cm^2^ at the furthest edges of the substrate (Figure 4A). Light was not detected outside the range of the extraction substrate (orange perimeter, Figure 4Aii), indicating reliable illumination of samples only, with the capacity for preparation of non-irradiated controls adjacent to the extraction substrate. No significant difference was observed when assessing irradiance output 3 cm or 0 cm from the substrate surface. Indicating a distance of 3cm was appropriate to acquire measurements of spectral and wavelength output (Figure 4C-D).

**Figure 3:**
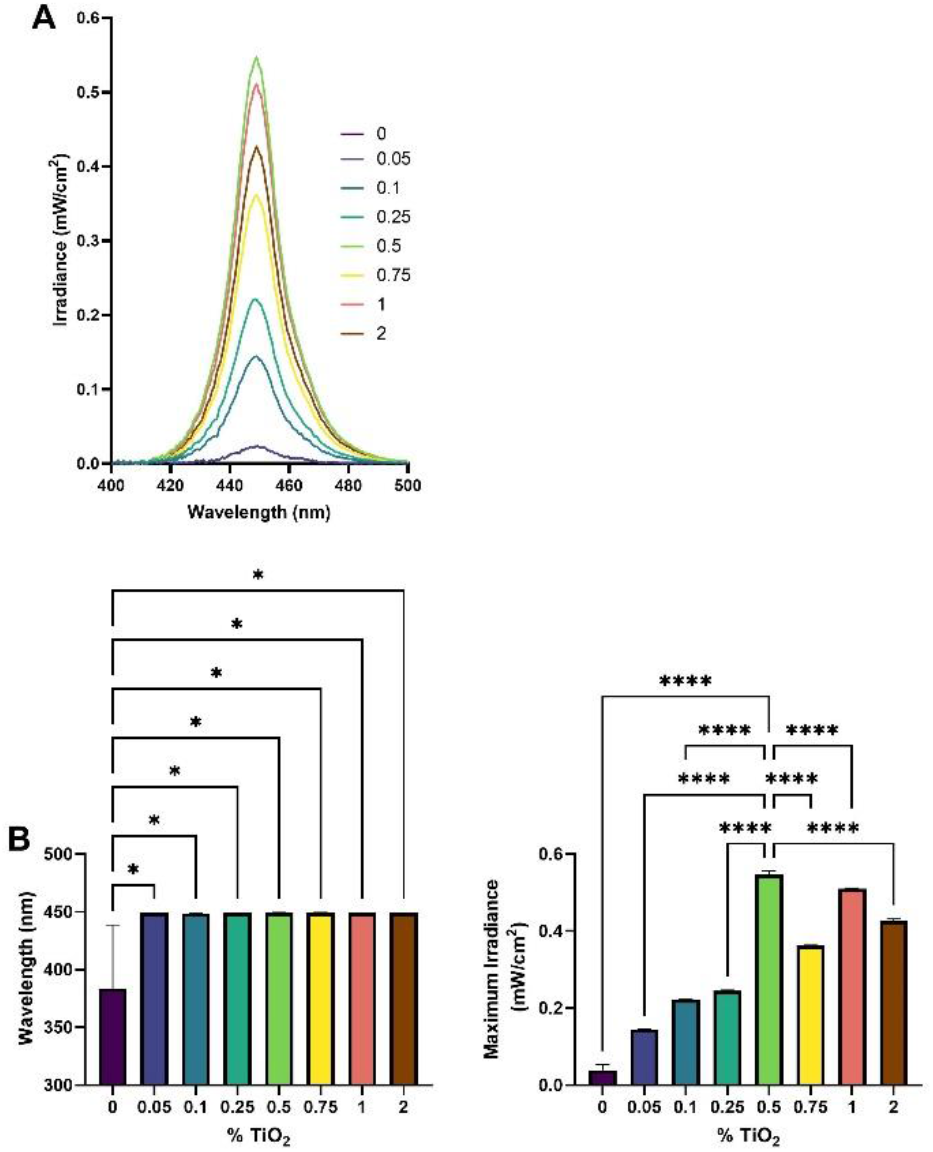
TiO_2_ potentiates irradiance output. Wavelength spectra (A), peak wavelength output (B) and maximum irradiance output (C) following deposition of various PDMS substrate comprising increasing concentrations of TiO_2_ were collected using a spectrometer coupled to Avantes software 3cm above substrate surface. Measurements recorded in triplicate and significance assessed via one-way ANOVA followed by Tukey test and denoted as ****P<0.0001, ***P<0.001, **P<0.01 and *P<0.05.

**Figure 4:**
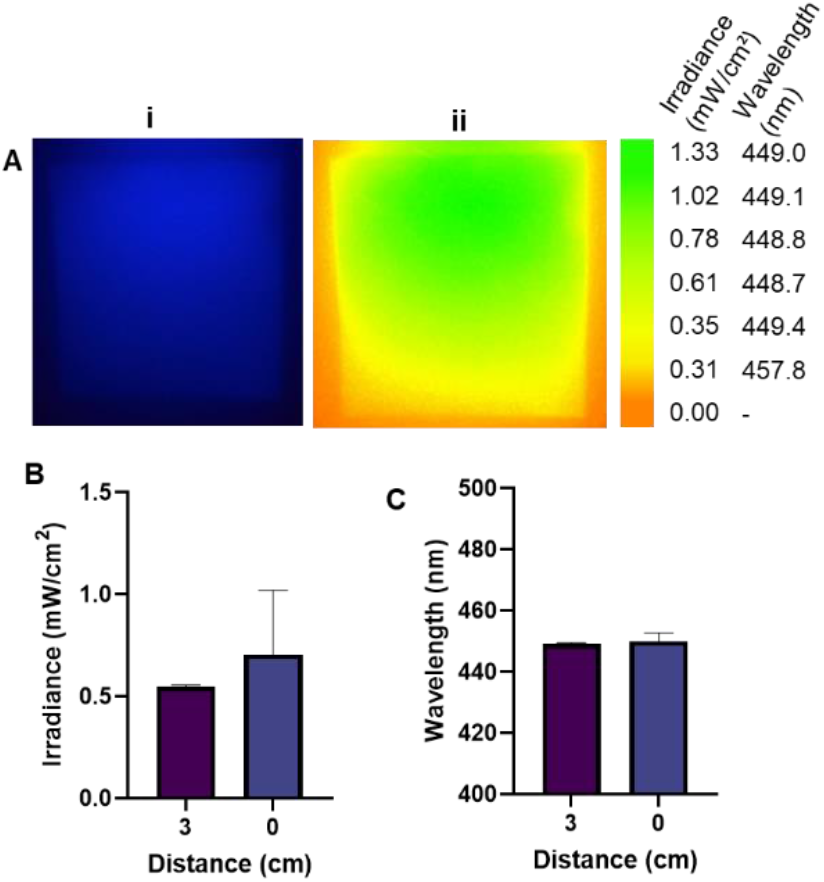
Irradiance output across the substrate surface. Irradiance maps acquired from calibrated CCD images (the μLED array is coupled at the top of these images) (i) and variability determined using image J (ii) of (A) 0.5% TiO_2_ substrate Irradiance (B) and wavelength output (C) measured in the centre of the platform using a spectrometer directly upon the substrate surface vs at a distance 3cm above the substrate. Measurements recorded in triplicate and significance assessed via unpaired t-test.

### Skin Commensals Diminish Irradiance Output from the 450 nm μLED illumination platform

Of the 16 species and strains surveyed, four diminished the irradiance signal output from the platform relative to the control but had no significant effect on wavelength output (Figure 5A, 6A). These included *M. luteus, K. arsenatis* and *K. rhizophila*, all of which display yellow pigmentation characteristic of carotenoid pigment expression and diminished irradiance output by 38.2% (p<0.01), 24.1% (p<0.05) and 37.3% (p<0.01) respectively relative to the glass control (Figure 5Ai). Measurement of the absorbance profiles of these species revealed triplicate peaks within the ranges of 416-422 nm, 440–457 nm and 465-485 nm (Figure 5Biv – vi). Indicating peaks particularly within the range of 440-457 nm could absorb light from the μLED platform. *S. hominis* also reduced irradiance output by 19.2% (p<0.05, Figure 6A) and displayed a peak in absorption profile of 420 – 422 nm (Figure 6Bv). *S. aureus* (Figure 6Bi), and *S. epidermidis* (Figure 6Biv) also exhibited spectral peaks within this wavelength range but had no significant effect on irradiance output. *S. capitis* and *S. hominis* also possessed a peak within the UV range (353 – 357 nm, Figure 6Biii and v).

**Figure 5:**
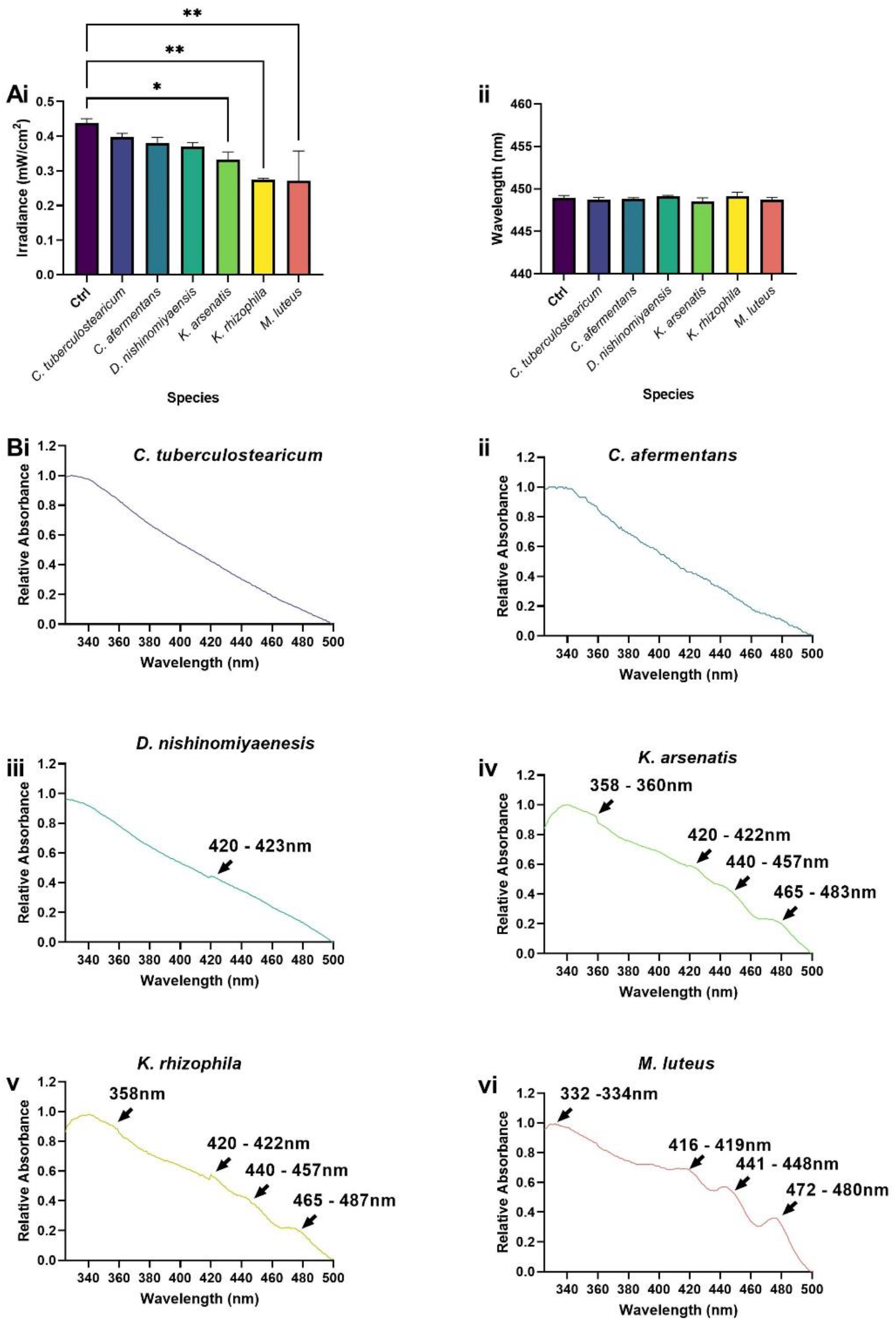
Diminished output of blue light is dependent upon skin commensal absorbance spectra. Skin commensals fixed to glass slides were applied to 0.5% TiO_2_ scatter substrate and irradiance (A) and wavelength output (B) were recorded. Absorbance spectra (C) of *C. tuberculostearicum* (i), *C. afermentans* (ii), *D. nishinomiyaenasis* (iii), *K. arsenatis* (iv), *K. rhizophila* (v) and *M. luteus* (vi). Measurements recorded in triplicate (N = 3) acquired relative to a glass control. Significance assessed via one-way ANOVA followed by Tukey test and denoted as ****P<0.0001, ***P<0.001, **P<0.01 and *P<0.05.

**Figure 6:**
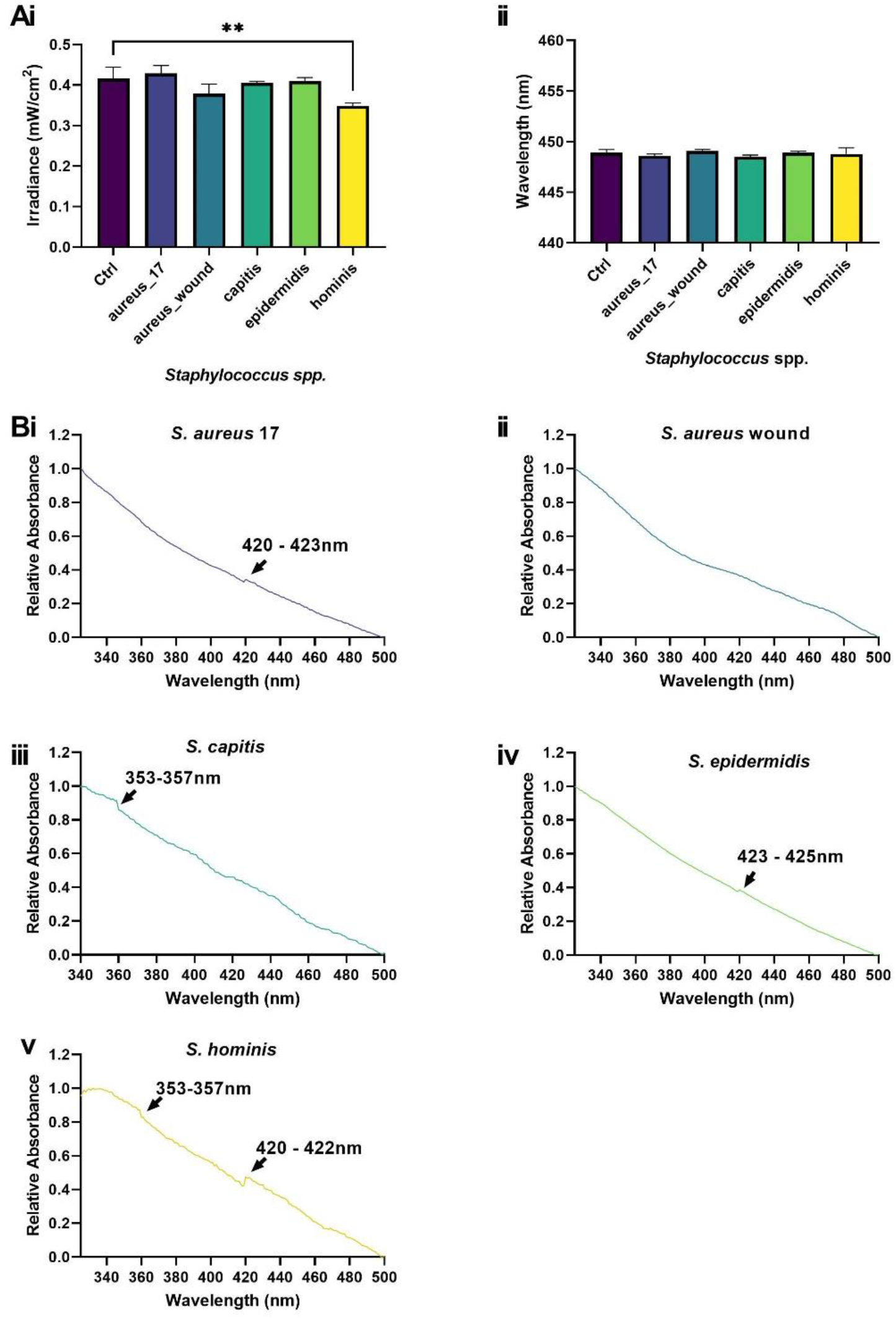
*Staphylococcus hominis* impacts blue light transmission. *Staphylococcus* spp. fixed to glass slides were applied to 0.5% TiO_2_ scatter substrate and changes in irradiance (A) and wavelength output (B) were recorded via spectrometry. Absorbance spectra (C) of *S. aureus* 17 (i), *S. aureus* wound (ii), *S. capitis* (iii), *S. epidermidis* (iv) and *S. hominis* (v). Measurements recorded in triplicate (N = 3) acquired relative to a glass control. Significance assessed via one-way ANOVA followed by Tukey test and denoted as ****P<0.0001, ***P<0.001, **P<0.01 and *P<0.05.

*Cutibacterium* spp. did not attenuate irradiance (Figure 7Ai), but *C. acnes* 177 and *C. granulosum* displayed absorbance spectra characteristic of porphyrin expression with peaks from 389 – 416 nm (Figure 7Bii-iii).

**Figure 7:**
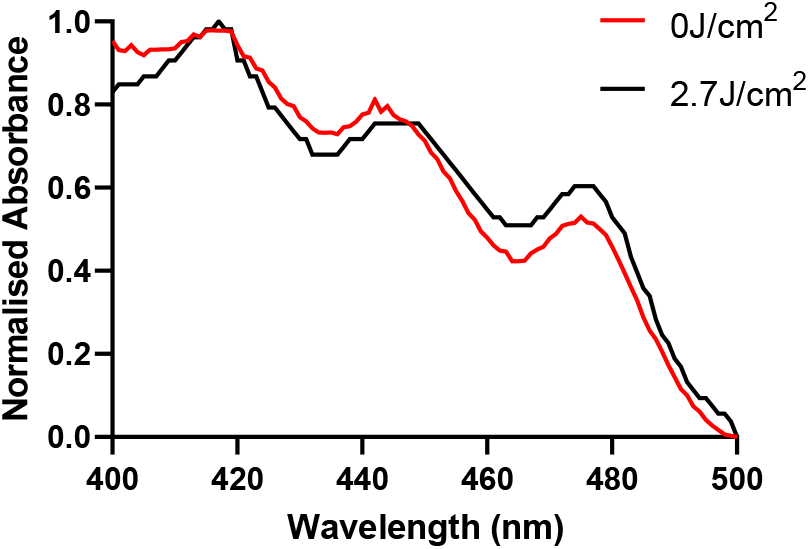
Exposure of *M. luteus* to blue light. Absorbance spectra of *M. luteus* pre– and post-exposure to blue light (n = 2, 450nm, 2.7J/cm^2^).

*M. luteus* exhibited a spectrum with defined peaks in the 450 nm range. Therefore, absorbance profiles were acquired pre and post-exposure to blue light (450 nm, 90 mins, 0.55 mW/cm^2^, and 2.7 J/cm^2^). Despite the low dose applied, a small shift in the absorbance profile was observed from 440 – 448 nm (peak; 444nm) to 442 – 449 nm (peak: 446nm), and elevated mean absorbance from 450 nm – 480 nm following irradiation (Figure 8).

**Figure 8:**
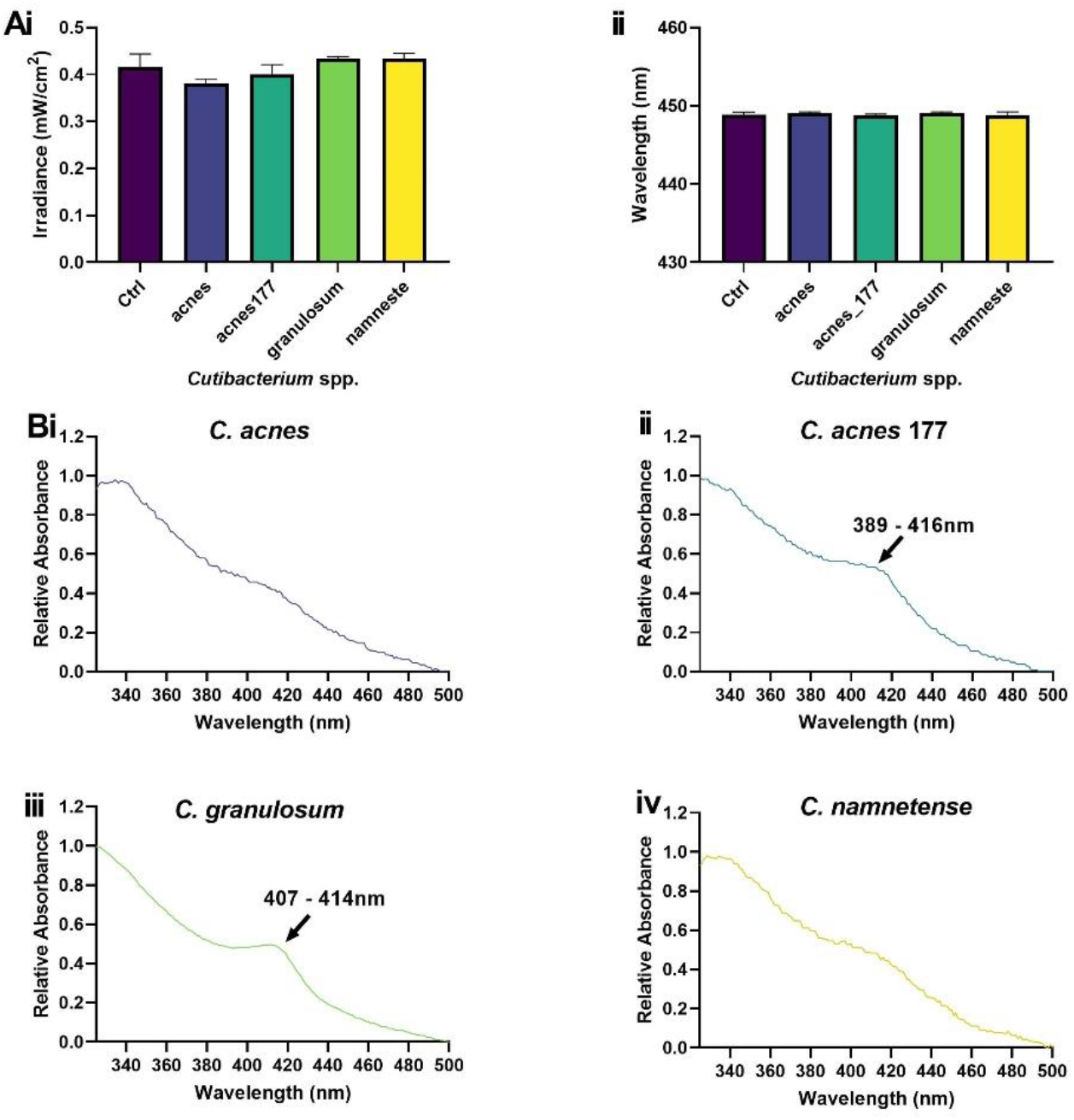
Selected *Cutibacterium* spp. exhibit peaks typical of porphyrin expression. *Cutibacterium* spp. fixed to glass slides were applied to 0.5% TiO_2_ scatter substrate and changes in irradiance (A) and wavelength output (B) were recorded via spectrometry. Absorbance spectra of *C. acnes* (i), *C. acnes* 177 (ii), *C. granulosum* (iii) and *C. namnetense* (iv). Measurements recorded in triplicate (N = 2) acquired relative to a glass control. Significance assessed via one-way ANOVA followed by Tukey test and denoted as ****P<0.0001, ***P<0.001, **P<0.01 and *P<0.05.

## 4 Discussion

Our bodies are home to a diverse and delicately balanced ecosystem. Tipping this balance can cause dysbiosis, resulting in reduced microbial diversity or overrepresentation of certain species within the community, characteristic of conditions including acne and atopic dermatitis (6, 33). Blue light (380 – 500 nm visible light) has high potential as a safe, non-invasive, and relatively cheap approach for the manipulation of microbial communities due to its microbicidal properties (34). However, studies employ poorly characterised light sources with variable intensity and spectral properties, resulting in conflicting findings (35). We present evidence of the thorough characterisation of a μLED illumination platform for *in vitro* studies. On this platform, substrates comprising 0.5% TiO_2_ disperse and enhance irradiance delivery to samples exploited in routine *in vitro* studies without impacting spectral output. This results in an average wavelength and maximum irradiance of 450 nm and approximately 0.5 mW/cm^2^ respectively (Figure 3). We note that the homogeneity of the irradiance could be further improved if desired by using a spatially graded concentration of TiO_2_ as we have previously shown (36).

The exploitation of this μLED platform ensures targeted light delivery, a feature often overlooked when ensuring the reliable assessment of the biological effects of light relative to a non-irradiated control (26, 37). Titanium dioxide (used as nanoparticles in the scatter membrane) exhibits a series of beneficial properties, not only does incorporation into PDMS provides potential for incorporation into a wearable flexible device suitable for clinical application (38), but also potentiates light delivery without increasing thermal output from a light source. Resulting in the application of a model system capable of discerning wavelength-specific effects on biological outputs, rather than potential undesired thermal effects (39). It also offers a dual advantage, in which photocatalysis of TiO_2_ has proven to induce the production of reactive oxygen species (ROS). This property may prove beneficial for potential decontamination applications (40).

Whilst a device emitting a wavelength of 450 nm was exploited in this study, blue light of 405nm is often regarded as the key wavelength in mediating microbicidal effects (41). This assumes that all microbial species are uniformly sensitive to the same single wavelength of blue light. However, responses to light are fundamentally dependent upon the presence or absence of wavelength-specific chromophores (14, 26). In this study, we demonstrate variability in absorbance spectra of bacterial species abundant across the acne, AD and healthy skin microbiomes (42). In which peaks in absorbance spectra denote the presence or absence of these wavelength-specific chromophores. Peaks around 422 nm proved particularly abundant across *Staphylococcus* and commensal species (Figure 5 – 7). We hypothesise these peaks could be attributed to the presence of components of the citric acid cycle (TCA) including flavin adenine dinucleotide or nicotinamide adenine dinucleotide, which exhibit autofluorescent properties and have been thoroughly characterised in bacterial species including *Pseudomonas aeruginosa* (43, 44). Staphylococci also possess a complete riboflavin synthesis system comprising a range of Flavin-containing complexes (45-47). Riboflavin exhibits an absorbance spectrum peak of 423 nm in water and the synthesis of riboflavin by *S. epidermidis* contributes to innate immune priming via the stimulation of mucosal-associated invariant T cells (48, 49). Along with *C. acnes, S. epidermidis* is cited as a key player in acne vulgaris pathogenesis and excitation with 405 nm light has proven to induce bacterial lysis via a reactive oxygen species-dependent mechanism (13, 50, 51). Collectively, the excitation of chromophores derived from both *C. acnes* and *S. epidermis* could contribute to the efficacy of blue light observed in the management of acne vulgaris clinically (52). Whilst commensal *Staphylococcus* spp. exhibited peaks within a range denoting potential Flavin expression there was no evidence of carotenoid expression from *S. aureus* strains used in this study, plausibly owing to their lack of pigmentation, a feature common in a large proportion of *S. aureus* strains (53).

The presence of a Soret band denoting porphyrin expression proved apparent in only one of the *C. acnes* strains surveyed. The capacity of *C. acnes* to express porphyrins has proven an important virulence factor and strains exhibiting higher levels of porphyrins (phylotype 1) are typically in higher abundance in individuals with acne vulgaris (54). Both *C. acnes* strains utilised were derived from healthy volunteers and belong to phylotype 1A1 (55). The strain of *C. granulosum* utilised in this study also proved to exhibit a strong peak denoting porphyrin expression (Figure 7). Previous studies have provided evidence that *C. granulosum* produces significantly higher levels of porphyrins relative to *C. acnes*, with particularly high expression of coproporphyrin III which exhibits a typical absorbance spectrum within the range of 400 – 415 nm (56, 57). Suggesting porphyrin expression from other *Cutibacterium* strains utilised may have been below the detection limit of the system used in this study.

We demonstrate species expressing carotenoid pigment peaks *(K. arsenatis, K. rhizophila* and *M. luteus*) diminish the irradiance output of our μLED platform by an average 30% (Figure 5, p<0.05). Indicating wavelength-dependent excitation of chromophores, and subsequent absorbance of light. To confirm this hypothesis, *M. luteus* was exposed to 450 nm light from the platform for 90 minutes and changes in absorbance pre– and post-exposure recorded. Despite the low dose applied (2.7 J/cm^2^), subtle shifts in absorbance spectra were observed from a peak at 440-448 nm pre-exposure to 442-449 nm post-exposure (Figure 8). Photobleaching of carotenoid pigments using blue light has been previously reported (58). However, how this impacts microbial physiology and in turn microbiome composition remains to be characterised. Current evidence suggests photodegradation of such pigments results in increased sensitivity to reactive oxygen species-dependent lysis (58), which in turn could trigger stress responses in certain species resulting in shifts in the expression of virulence and biofilm formation associated genes (59).

In conclusion, this study demonstrates evidence of a robust μLED illumination platform the design of which could prove highly applicable to in-vitro photobiomodulation studies but also to management of skin conditions. We show that members of the skin microbiome are capable of manipulating spectral output from this device and this mapped to the presence of wavelength-dependent chromophores. Based upon these findings, we can now take strides towards mapping how the expression of chromophores might impact microbial responses to blue light and in turn how we can exploit these findings for the management of prevalent skin conditions.

## 5 Back Matter

### Funding

Financial support for this project was provided by the University of Manchester Research Collaboration Fund (P127221) and The Leverhulme Trust Research Leadership Grant (RL-19-038).

#### Acknowledgements

We thank Xiangyu He for assistance in the fabrication of the μLED and Dr Faye adel a Aldehalan for the provision of bacterial strains used in this study. We also like to thank the University of Manchester Genomic Technology Core Facility for 16s sequencing of bacterial strains.

## Data Availability Status

Data underlying the results presented in this paper are available via the following DOI: 10.48420/25316077.

